# 3D RNA from evolutionary couplings

**DOI:** 10.1101/028456

**Authors:** Caleb Weinreb, Torsten Gross, Chris Sander, Debora S. Marks

## Abstract

Non-protein-coding RNAs are ubiquitous in cell physiology, with a diverse repertoire of known functions. In fact, the majority of the eukaryotic genome does not code for proteins, and thousands of conserved long non-protein-coding RNAs of currently unkown function have been identified. When available, knowledge of their 3D structure is very helpful in elucidating the function of these RNAs. However, despite some outstanding structure elucidation of RNAs using X-ray crystallography, NMR and cryoEM, learning RNA 3D structures remains low-throughput. RNA structure prediction *in silico* is a promising alternative approach and works well for double-helical stems, but full 3D structure determination requires tertiary contacts outside of secondary structures that are difficult to infer from sequence information. Here, based only on information from RNA multiple sequence alignments, we use a global statistical sequence probability model of co-variation in a pairs of nucleotide positions to detect 3D contacts, in analogy to recently developed breakthrough methods for computational protein folding. In blinded tests on 22 known RNA structures ranging in size from 65 to 1800 nucleotides, the predicted contacts matched physical nucleotide interactions with 65-95% true positive prediction accuracy. Importantly, we infer many long-range tertiary contacts, including non-Watson-Crick interactions, where secondary structure elements assemble in 3D. When used as restraints in molecular dynamics simulations, the inferred contacts improve RNA 3D structure prediction to a coordinate error as low as 6 – 10 Å rmsd deviation in atom positions, with potential for further refinement by molecular dynamics. These contacts include functionally important interactions, such as those that distinguish the active and inactive conformations of four riboswitches. In blind prediction mode, we present evolutionary couplings suitable for folding simulations for 180 RNAs of unknown structure, available at https://marks.hms.harvard.edu/ev_rna/. We anticipate that this approach can help shed light on the structure and function of non-protein-coding RNAs as well as 3D-structured mRNAs.

## Introduction

Structured RNAs play diverse cellular roles, including the regulation of mRNA splicing, degradation and localization^1-5^, delivery of viral genome^6^, X-inactivation^7^, as well as its more well known roles in translation and ribosomal processing^8^ This probably represents only a fraction of RNA’s functional repertoire. High-throughput approaches have revealed pervasive transcription throughout the non-coding genome. Although much of this transcription may be artifactual or biological noise^9^, it has nevertheless revealed some functional RNAs, including long noncoding RNAs^10^ that may have three-dimensional structures that can act can act – for example – as protein scaffolds^11^. Other large-scale evidence of structure within the coding region of mRNAs has emerged from new high-throughput approaches that detect base-pairing *in vivo*^12-15^.

Many of these newly discovered RNAs act by adopting specific 3D structures^16,17^ but high-resolution structure determination remains labor intensive. Thus, there is a renewed interest in computational approaches to structure prediction. Though 3D structure prediction for smaller RNAs has become moderately successful ^18-22^, predicting the structure of large (> 70 nt) RNAs without experimental information remains challenging^23^ as the RNAs with multiple helical segments can assume diverse folds, producing a conformational search space that is impossibly large. Information about contacts between bases that are distant in the secondary structure but close in the 3D structure (tertiary contacts) substantially narrows the conformational search space^24^. Biochemical probing studies can provide evidence for low-resolution tertiary contacts^25,26^, and 3D structures computed using extensive experimental chemical mapping information have achieved accuracy of ~ 8Å positional root mean square deviation (rmsd) from the known crystal structures. However, predicting RNA 3D structure completely *in silico* has been challenging^27^.

The recent explosion of available homologous sequences for RNAs provides an exciting opportunity to detect tertiary contacts from sequence co-variation alone. However, existing methods – designed to detect local (pairwise) patterns of sequence co-variation – have had limited success^28,29,30^. Since RNA tertiary contacts often form complex networks^31^, it is possible that overlapping patterns of sequence constraint interfere with each other, obscuring true correlations and producing spurious transitive correlations when multiple contacts are chained together. A similar problem stymied protein structure prediction until the emergence of global maximum entropy models that could de-convolve the underlying network of residue-residue interactions^32-34^. Although local, non-global co-variation models have successfully predicted RNA secondary structure, we hypothesized that global maximum entropy models such as those used in predicting protein 3D structure^35-38^ would enhance prediction of 3D contacts in RNA, including tertiary contacts such as non-WC base pairs. Here we adapted the maxent model to RNA sequences alignments and tested the ability of the approach to predict tertiary structure contacts on 22 known RNA structures. In genuine prediction mode, we infer 3D contacts for 180 known functional RNAs with no known 3D structure from mulitple sequence alignments alone.

## Results

### Evolutionary couplings accurately predict 3D contacts

We adapted the evolutionary couplings model used for proteins to predict ECs for RNA and evaluated the ability of ECs to recover RNA 3D contacts for a specific RNA from each of the 22 RFAM families that had a known 3D structure (Supplementary Table 1). To assess the accuracy of predicted contacts, we defined contacts as true positives when their minimumatom-distance was < 8 Å. All 22 RNAs had a true positive rate above 70% for L/2 ECs (where L is the number of nucleotides) and 8 had over 80%. ECs predicted contacts with greater accuracy than mutual information (MI), which has been widely used for RNA secondary structure prediction^39,40^ (Fig. 1A). This was true even when using enhanced MI (MIe) that implements two features of the EC statistical model: (1) Down-weighting of similar sequences to avoid spurious correlations from phylogeny; (2) An average product correction (APC)^41^. We refer to MI without these modifications as raw MI (MI_R_).

**Figure 1:**
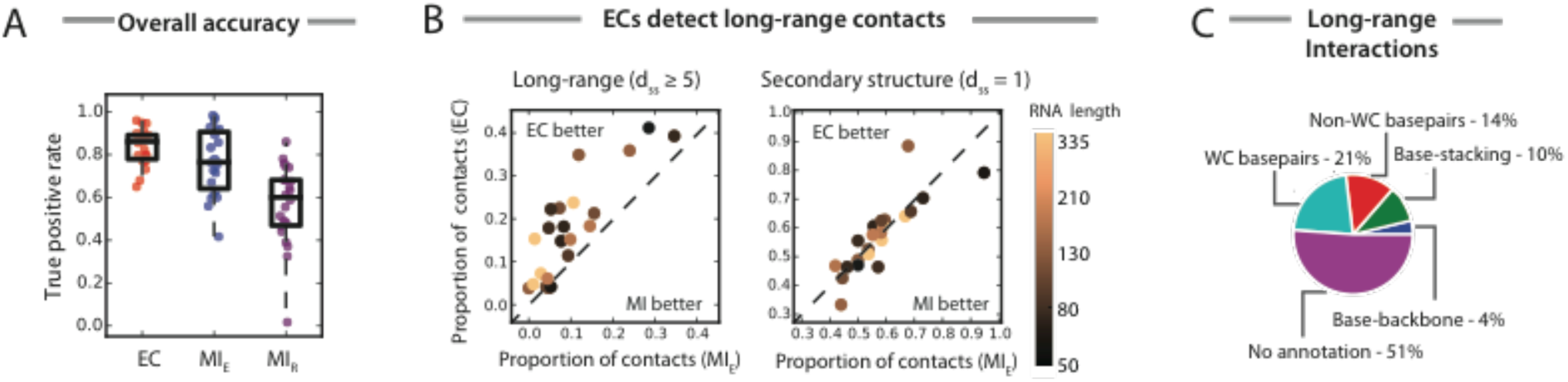
Comparison of EC to MI: Summary of 22 alignments. (A) EC predicts structural contacts with an overall higher accuracy than MI_E_ or MI_R_. To see if EC and MI are sensitive to different types of contacts, we classified contacts based on their distance in the secondary structure and defined three categories: secondary structure contacts (d_ss_ = 1); short-range contacts (d_ss_ < 5); and long-range contacts (d_ss_ ≥ 5). (B) Whereas EC and MI detect a similar number of secondary structure contacts, ECs are significantly enriched with long-range contacts. (C) The long-range contacts detected by ECs represent a variety of biochemical interactions, as annotated from the crystal structure. Notably, 51% of these contacts have no annotation. Understanding the biochemical basis for these interactions is a promising area for future research.

### ECs detect long-range interactions

Though overall accuracy is important – not all contacts are created equal. Often, complex RNA folds are stabilized by a small number of critical long-range contacts that bridge distant parts of the secondary structure. Detecting these contacts computationally would be a significant advance since they provide crucial structural information that cannot be inferred from the secondary structure alone. To test whether ECs could detect long-range contacts, we define the secondary-structure-distance (*d_ss_*) between two bases as the length of the shortest path between them in a graph where nodes are bases and edges are either WC base pairs of instances of adjacency on the RNA chain. True positive predicted contacts can then be divided into three categories: WC base pairs (*d_ss_* = 1); short-range contacts (1 < *d_ss_* < 5); and long-range contacts (*d_ss_ ≥* 5). Compared to MI_E_, EC-predicted contacts are highly enriched with long-range contacts (0.66 – 8 fold; mean fold-change = 2.4; *p* ≤ 10^−3^; Fig. 1B). Indeed, we were able to robustly predict long-range contacts across the 22 alignments, obtaining on average 0.05 * *L* contacts for an RNA of length *L,* with some notable outliers. For example, for Ribonuclease P, a ribozyme of length *L* = 334, we correctly predicted 53 long-range contacts using ECs, compared to 9 for MI_E_ and 10 for MI_R_. The long-range contacts predicted by ECs represent a variety of biochemical interactions, including base pairing, base stacking and base-backbone interactions (Fig. 1C). Remarkably, 51% of long-range contacts had no annotation in the crystal structure. These contacts may arise from un-annotated interactions such cooperative binding of an ion, ligand, or protein, or through patterns of hydrogen bonding that are not currently characterized. Understanding the biochemical basis for these interactions is a interesting area for future research.

### ECs detect non-WC base pairs

Pairs of RNA bases often form contacts through hydrogen bonding, assuming geometrical configurations that can broadly be divided into Watson-Crick (WC) base pairs and non-WC base pairs. Previous work has suggested that non-WC base pairs cannot be predicted from sequence co-variation alone^42^, since they do not sufficiently coevolve. Others have hypothesized that non-WC base pairs cannot be predicted computationally since they often participate in interaction networks, with overlapping patterns of sequence constraint obscuring the underlying pairwise co-variation^31^. Evolutionary couplings derived from global maximum entropy models have successfully deconvolved similar networks of constraints within proteins, and may be useful in this context too. We found that ECs detect non-WC base pairs with unprecedented sensitivity, detecting substantially more pairs than MI_E_ (0.5 – 8.5 fold; mean fold-change = 1.92; *p* ≤ 10^−3^). Overall, ECs captured 16% of all annotated non-WC base pairs across the 22 structures, which may represent a high level of sensitivity, since not all non-WC base pairs coevolve. We expect that ECs will complement existing approaches for detecting non-WC base pairs that rely on the concept of isostericity^30^, since they can focus attention on interactions with the strongest co-evolutionary signal.

### ECs reveal contacts in the eukaryotic ribosome

ECs may be sensitive to interacting nucleotides in large RNAs that form topologically complex folds with abundant long-range contacts. ECs computed on a full alignment of eukaryotic ribosomal sequences (RF01960) are over 90% accurate for the top 900 (L/2) contacts and predict substantially more WC and non-WC base pairs than MI_E_ (Fig. 2A-C). ECs also predict many long-range contacts between nucleotides (Fig. 2D). However, some contacting nucleotides are not predicted, including a large pseudo-knot between the 5-prime and 3-prime regions. ECs may not capture these contacts due to a lack of sensitivity. Alternatively, these regions may not actually be co-evolving and their proximities could be a simple consequence of the geometric constraints from the remainder of the molecule. Distinguishing contacts that genuinely coevolve from those that form due to geometrical constraints is important because coevolving contacts are conserved across the alignment and likely to be functionally important.

**Figure 2:**
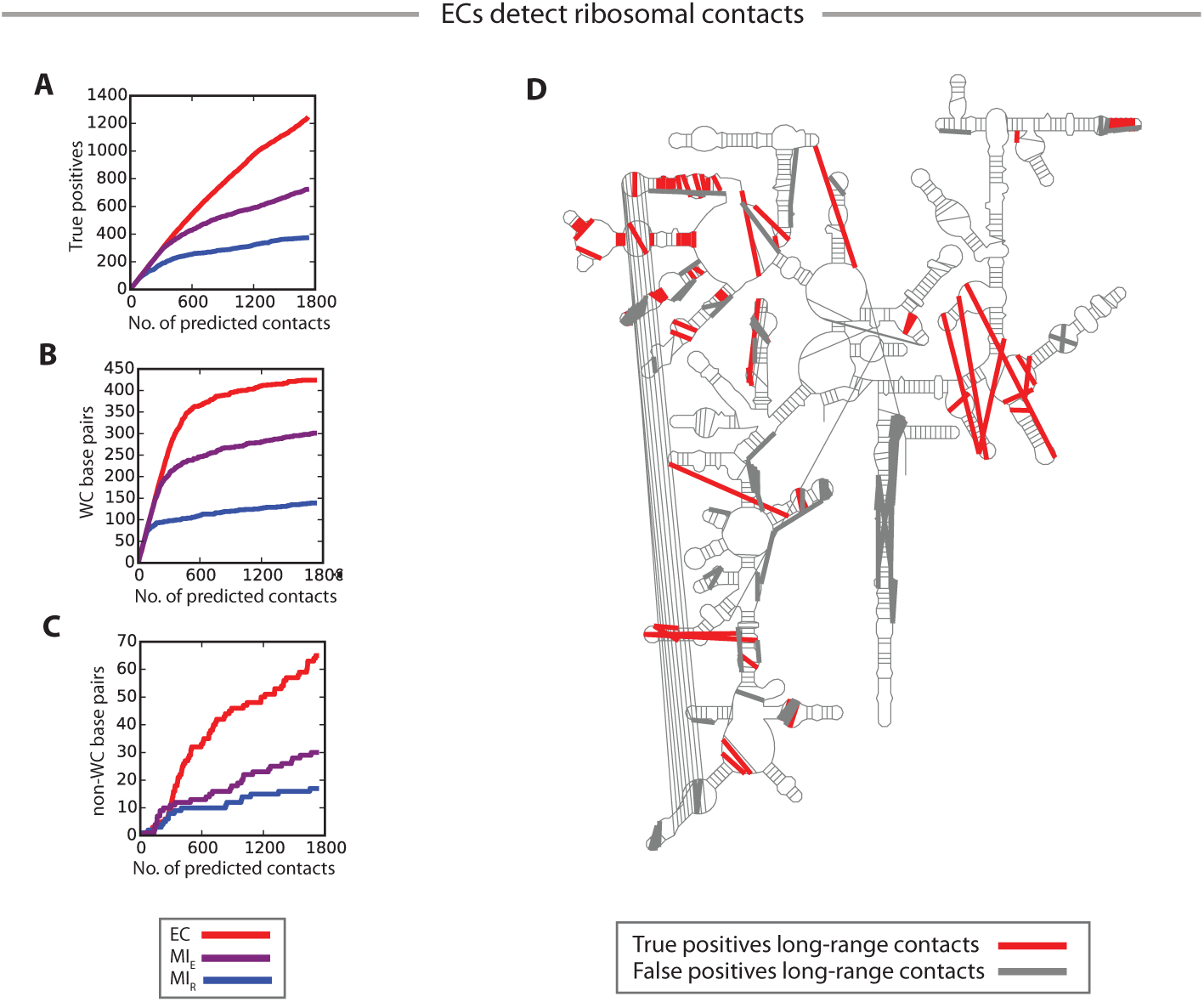
ECs detect ribosomal contacts. ECs may be sensitive for large RNAs that form topologically complex folds with abundant long-range contacts. We tested this notion on the 40S ribosome (RF01960), and found a dramatic difference between ECs and MI, with ECs detecting more overall true positives (A), more Watson-Crick base pairs (B) and more non-WC base pairs (C). ECs also detect ~50 long-range contacts that bridge distance parts of the secondary structure (D).

### Evolutionary couplings reveal functionally important contacts in riboswitches

To probe whether evolutionarily coupled tertiary contacts are functionally important, we investigated the top scoring ECs in riboswitches, which are cis-acting regulatory RNA genes that undergo ligand or temperature^45^ dependent conformational changes between at least two mutually exclusive functional states^43-45^. In four riboswitches from our dataset – *S*-adenosylmethionine (SAM), active vitamin B_1_ (TPP), active folate vitamin (THF), and Adenine-sensing ( Purine) – we found a cluster of tertiary contacts that stabilize the ligand bound conformation, but are broken in the unbound conformation (Fig.3). For instance, in the TPP riboswitch, ECs between the L5 loop and J3-2 helix (69A-37G, 69A-23C and 70A-22C, in numbering from 2gdi.pdb^46^) have stacking interactions when the TPP ligand binds^46^, Fig. 3B. The EC predicted contacts that are part of the ligand bound conformations are not present in the secondary structure or detected by MI_E_. Similarly, very high ranking ECs between nucleotides between P2-P4 and P1-P4 in the SAM riboswitch form conditionally on binding of *S*-adenosylmethionine, (C25-G89, U26-A88, G27-C87, G28-C86, A9-A84, A10-A84 in numbering from 4kqy.pdb^47^). Some of these EC predicted contacts are also detected by the MI_E_ but there is a cluster between P1-P4 that are not, including base stacking between A9-A84. In addition to tertiary contacts from the known ligand-bound conformation, ECs could be used to confirm the existence of hypothesized contacts in the unbound conformation, which has no known structure. Unfortunately, many of these contacts involve nucleotides that are not currently represented in the RFAM family. We look forward to analyzing these more complete alignments.

**Figure 3:**
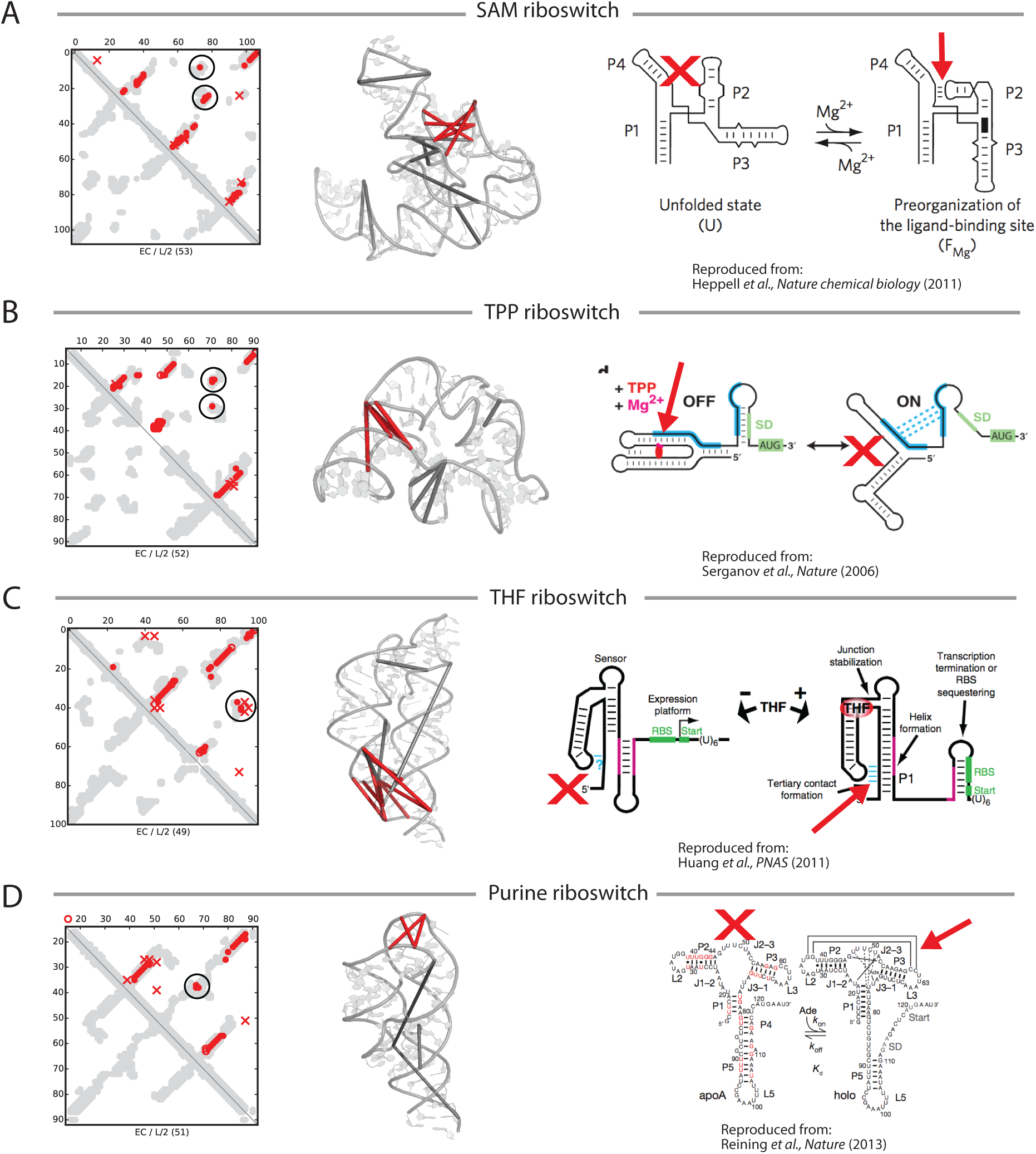
ECs identify functional interactions in riboswitches. 3D contacts detected through using ECs are conserved across the RNA family, and may therefore be functionally important. In four riboswitches, the most significant tertiary interactions revealed by ECs are critical for stabilizing the ligand-bound conformation. In each example, a contact map (left) shows the top L/2 contacts. The circled contacts – which are highlighted red on the 3D structures (middle) – are formed in the ligand-bound state, but violated in the unbound state. This is illustrated by the schematics (right), which were reproduced from prior studies.

### Proof of principle-folding

To understand the function of non-coding RNAs, it is critical to know their structure. Currently, a major obstacle in RNA 3D structure prediction is the limited knowledge of long-range contacts that bridge distant parts of the secondary structure. We hypothesized that ECs could provide the critical tertiary contacts that are necessary to fold RNAs. Using coarse-grained molecular dynamics implemented in NAST^31^ followed by simulated annealing with XPLOR^48^, we predicted the all-atom structure for four RNA families (Fig. 4). Our predicted structures had an all atom RMSD of 6 – 12 Å, which is comparable to the state of the art for RNA structure predictions that have tertiary contacts derived from biochemical probing^23^.

**Figure 4:**
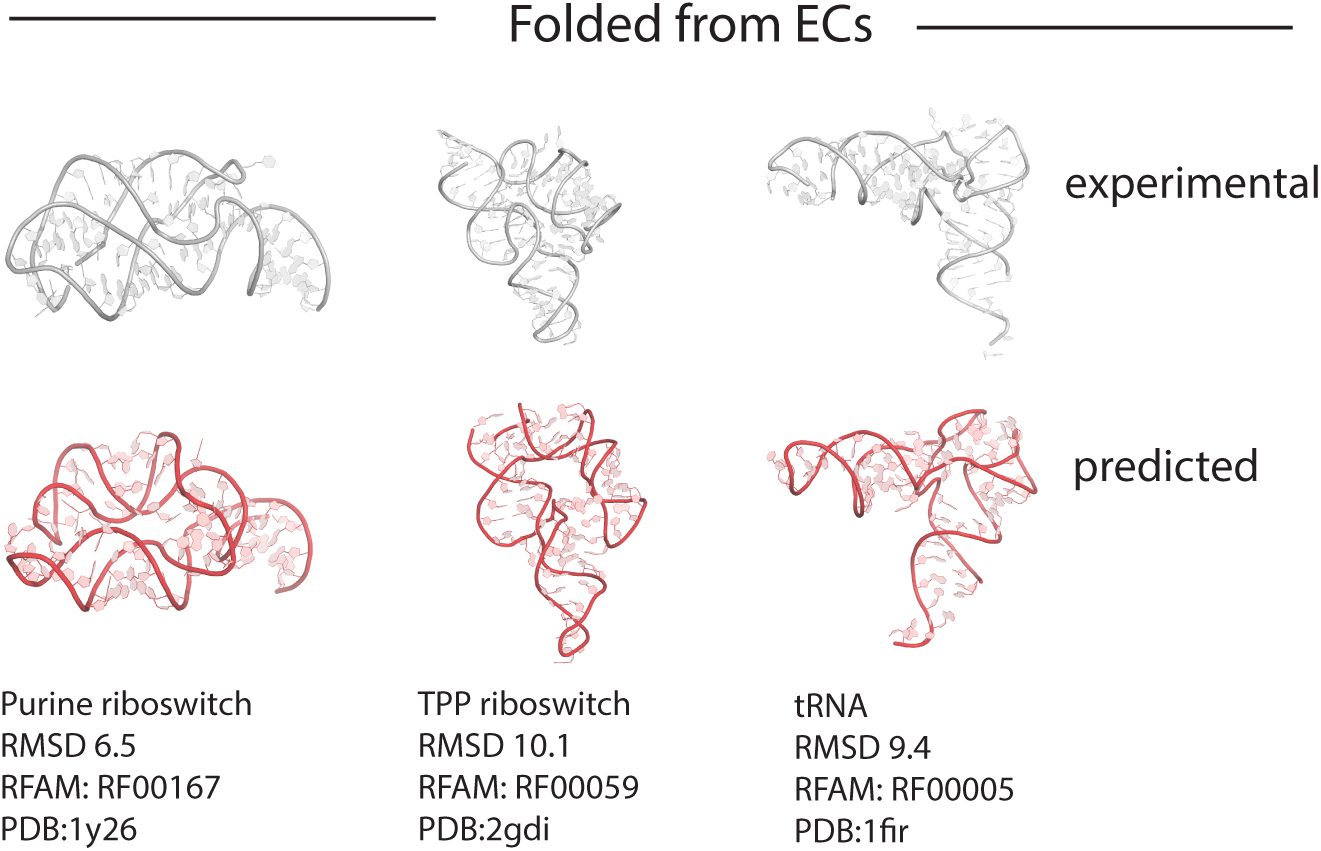
Blinded structure prediction. EC contacts provide enough structural information to predict the 3D structure of medium sized (70 – 120 nt) RNAs. The results from four predictions are shown here, with the true structure shown above (gray) and the predicted structure shown below (red).

### Contact Predictions of RNAs of unknown 3D structure

We predicted the 3D contacts for 160 RNA genes represented in RFAM that have no 3D structure known for any member of the family. This includes members of the Group-II catalytic introns whose 3-dimensional arrangement has been elusive since their discovery over 25 years ago^49^ Not surprisingly, the high ECs (red dots, Fig. 5A) capture the predicted secondary structure (green dots, Fig. 5A) but they also capture clusters of contacts that connect together the predicted helices into a more compact structure (black circles, Fig5A). Specifically, the EC pairs connect the end of stem loop 1 with the start of stem loop 3, end of stem loop 2 with end of stem loop 4 etc. Similar patterns in all the Group II intron predictions suggest compact globular structures that are independently known to exist. ECs for the small nucleolar RNA 25 connect the two prominent stem loops, predicting a compact folded RNA (Fig. 5B right panel). The prediction of the archeal RNAseP (left panel, Fig.5B) shows a similar 3D arrangement of the stem loops to those known for the other RNaseP families, though they are not sequences related enough to be put in same RFAM family, they are grouped into the same clan and their similar function suggests that the prediction is correct. Similarly, ECs of the SAM-IV riboswitch are similar to known functionally related SAM riboswitches, indirectly supporting the predicted 3D contacts (Fig. 5B middle panel).

**Figure 5:**
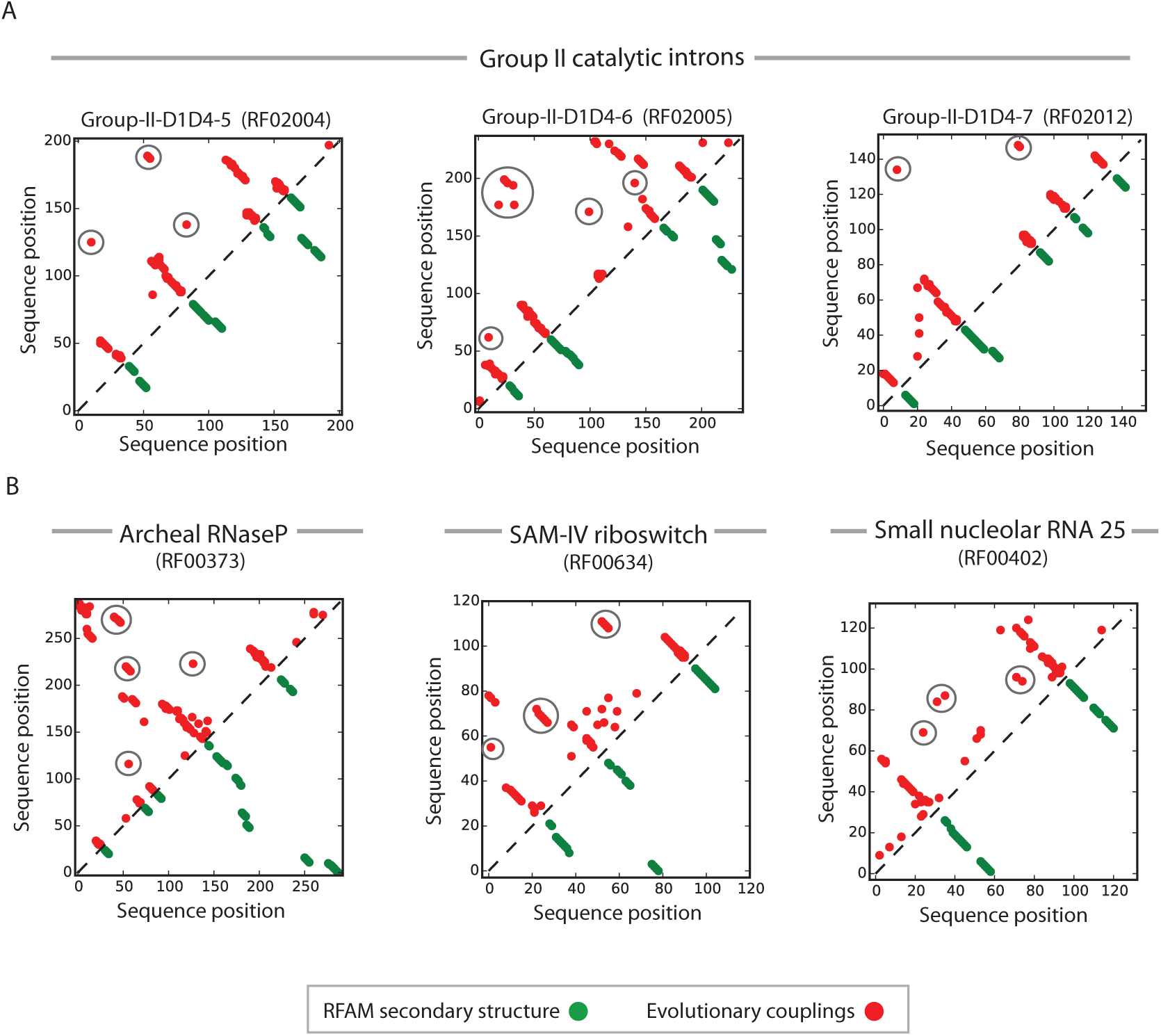
Predicted contacts for six RNA families. We use ECs to predict contacts for 160 RNA families with no known structure. Six families are presented here, selected for the presence of interesting tertiary contacts, which are highlighted with black circles. Many of these contacts appear to bridge the loops at the end of hairpins, and may be informative for 3D structure prediction.

Many more riboswitches are thought to exist than are currently known, but detecting them computationally has been challenging as they are often distant in primary sequence though machine learning based approaches represent a promising advance^50,51^. Here, we have discovered a conserved co-evolutionary motif among riboswitches in which tertiary contacts between stem-loops stabilize the ligand bound conformation and are broken in the unbound conformation. Searching for this pattern of evolutionary couplings could complement existing approaches for riboswitch detection.

## Discussion

We show that the global statistical models used for protein contact prediction give more accurate contacts than standard approaches and more importantly reveal nucleotide contacts that are important for the RNA function. In particular, the inferred evolutionary couplings (ECs) reveal clusters of contacts between loops and helices that are strategically located for the formation of 3D structures. In many cases the co-evolutionary signal from these interactions is drowned out by transitivity when using a purely local correlation model such as mutual information (MI), which is independently calculated for each pair of positions.

This work uses RNA alignments available in the RFAM database and we expect that the information content of the alignments could be improved with algorithmic refinements, especially by careful extension beyond the boundaries of current RFAM domains. Fortunately, ECs for RNA can be calculated with far fewer sequences than for proteins, since the number of parameters in the model scales with the square of the number of residue states, which is just four for RNA as opposed to twenty.

RNA secondary structure, which is formed by strings of hydrogen-bonded base pair interactions, can be well predicted by a variety of methods, including local co-variation methods such as MI. However, RNA gene products can form other kinds of inter-nucleotide contacts that are important for its 3D structure and function. With such non-secondary structure contacts derived from maximum entropy co-variation analysis, we perform proof-of-principle folding that uses only ECs and secondary structure prediction from RFAM. The resulting structures in some cases are better than those computed using detailed experimental constraints, even without the use of fragments or canonical helical constructs.

For practical reasons, we have focused attention on small and medium size RNA species in the range of tens to a few hundred nucleotides in length. For the largest molecule analyzed here, the small ribosomal subunit with about 1800 nucleotides, we derived predicted contacts, outside of secondary structure helices, at a 2:1 correct:false ratio, with broad distribution over the known tertiary structure contacts. However, complete folding of this and the large subunit of the ribosome probably requires assistance of ribosomal proteins. Evolutionary coupling analysis between RNA and proteins is of interest in this context and beyond the scope of this report.

The inferred evolutionary couplings, for each pair of residue positions, have a numerical value reflecting the strength of the interaction that the particular pair contributes to the mutational correlations observed for all pairs. Plausibly, the extent of translational propagation of the basic direct interactions in the entire system is stronger in proteins than in RNA molecules. We were therefore surprised that the maximum entropy method for extracting causative interactions works well in about one third of the currently analyzed sequence-rich RNA families and leads to reasonably accurate predictions of 3D tertiary contacts and 3D atomic structures. With the rapid acquisition of genetic sequences, we expect many more functional RNA molecules to reveal interesting 3D structures via maximum entropy contact analysis.

## Methods

### Selection of RFAM families

To test whether ECs could predict tertiary structure contacts, we used RNA multiple sequence alignments from the RFAM 11.0 database^52^, removing columns with > 50% gaps. We restricted to families where the effective number of sequences (Meff, see below) was greater than 0.5L, where L is the number of columns in the alignment, yielding 244 families (see Supplementary Table 1). Of these, 21 aligned to a known structure in the PDB ^53^. Data on these 21 structures are presented in Figure 1. For blind structure prediction, we removed structures with a length > 200 nt, as well as structures where the RNA of interest is part of a larger complex. Data on the remaining 13 structures are presented in Figure 3.

### Computing ECs

We applied a maximum entropy model to identify evolutionarily coupled pairs of columns in the alignments as described previously^36^. We inferred the parameters of our model using penalized Maximum Likelihood with a pseudo-likelihood approximation (pseudo-likelihood maximization; PLM)^36,54-57^ rather than with a previously applied mean-field approximation^32,33,35,37^. This method assumes that sequences are independent draws from an underlying distribution over sequence space. However, this assumption does not hold in reality, since many sequences are related by phylogeny. To account for this, we reweighted sequences in inverse proportion to their number over 80% similar neighbors. The sum of the resulting weights represents the effective number sequences (M_eff_).

For regularization, we used an L2 penalty with λ_h_ = 0.01 for single column fields and λ_e_ = 20.0 for the pair couplings. We also applied an average product correction (APC)^41^ to account for differences in the entropy of each column.

### Computing MI

To investigate how ECs compare to previous measures of co-evolution, we computed two versions of mutual information (MI). First we computed the raw MI (MI_R_) as shown below, where *f_i_*(*A*) = *P*(*S_i_* = *A*) and *f_ij_*(*A, B*) = *P*(*S_i_* = *A, S_j_* = *B*) for a sequence *S* in the alignment.

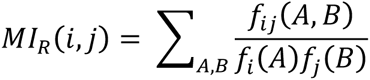

EC scores differ from MI_R_ in three ways: (1) They rely on a global maximum entropy model; (2) They down-weight sequences with a greater phylogenetic representation in the alignment; (3) They include an APC correction. Since feature (1) is the focus of this study, we also computed an enhanced MI score (MI_E_), which incorporates features (2) and (3), as has been done in previous work on RNA co-evolution^41^.

### Annotating interactions

For each alignment, we investigated the top L/2 contacts with a chain-distance > 4. We first classified contacts as true-positives if the minimum-atom-distance from the crystal structure was < 8 Å These were classified according secondary structure distance (d_ss_) and biochemical interaction type. The d_ss_ for a pair of bases is the length of the shortest path between them in a graph where nodes are bases and edges are either secondary-structure contacts or instances of adjacency on the chain. To compute d_ss_, we used the consensus secondary-structure provided by RFAM, which is inferred using a profile stochastic context-free grammar^58^. To classify contacts create random unfolded by their biochemical interaction type, we used crystal structure annotations from FR3D^31^ which were downloaded from RNA3DHub (http://rna.bgsu.edu/rna3dhub/).

### 3D structure prediction

We performed blind structure prediction using the Nucleic Acid Simulation Tool (NAST)^59^. NAST is a coarse-grained modeling tool that uses a combination secondary structure and tertiary contacts as inputs. For each RNA family, we generated 200 random unfolded structures that satisfied the secondary structure constraints (Fig. 6A). Next, we performed molecular dynamics using tertiary structure restraints to generate candidate models (Fig. 6B). To obtain these tertiary structure restraints, we used the ¾*L contacts with the top EC scores, where L represents the length of the RNA. Since the resulting lists contained many intra-helical contacts that are not useful for folding, we removed all contacts with d_ss_ < 5. We next sought to eliminate tertiary structure restraints derived from false-positive contacts. To that end, we performed molecular dynamics using weak constraints to iteratively remove restraints that were consistently violated by the resulting structures (Fig. 6C), removing at most 15% of the using simulated annealing contacts in any one round. Contacts were defined to be violated when the average distance between the corresponding bases was > pipeline 15 Å. Once all contacts were satisfied, we clustered the 20% of structures with the lowest NAST energy (Fig. 6D) using Biopython, producing n = 4 clusters. From each cluster, we chose a representative with the lowest NAST energy and then created an all-atom structure by assembling fragments from the ribosome (Fig. 6E). Finally, we refined the all-atom models by simulated annealing with XPLOR (Fig. 6F).

**Figure 6:**
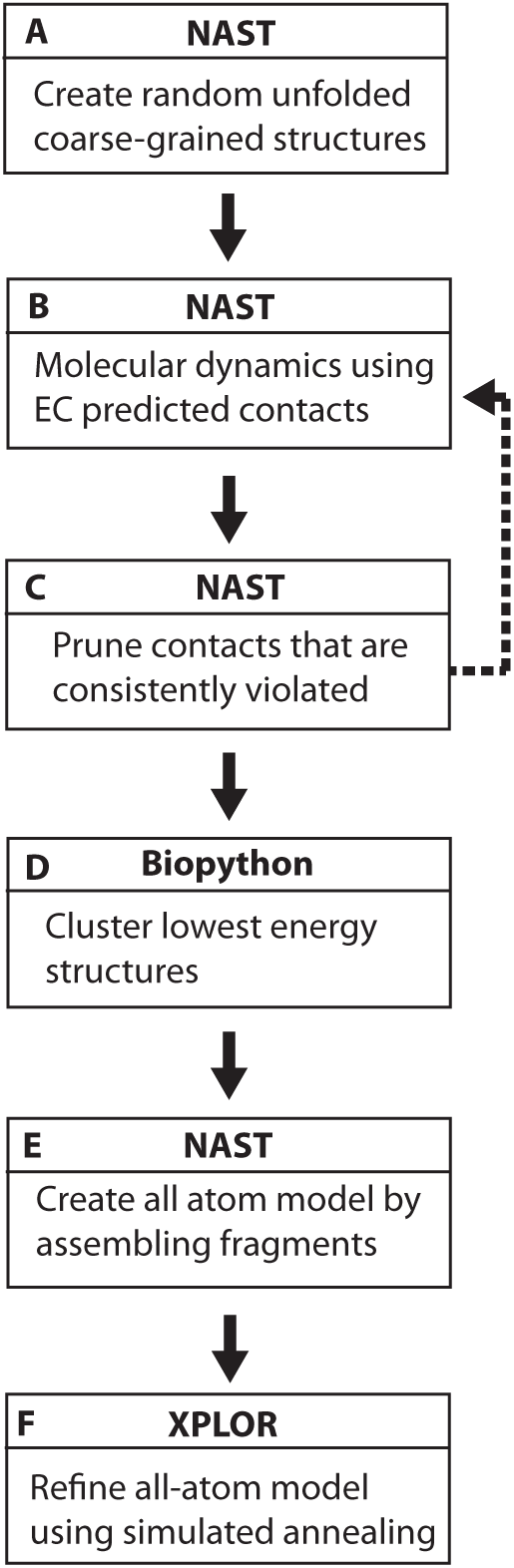
Overview of folding pipeline.

